# Establishment and Characterization of a Multi-Purpose Large Animal Exposure Chamber for Investigating Health Effects

**DOI:** 10.1101/415604

**Authors:** Xinze Peng, Mia R. Maltz, Jon K. Botthoff, Emma L. Aronson, Tara M. Nordgren, David D. Lo, David R. Cocker

**Affiliations:** Department of Chemical and Environmental Engineering, Bourns College of Engineering, Center for Environmental Research and Technology, University of California Riverside; BREATHE (Bridging Regional Ecology, Aerosolized Toxins, & Health Effects) Center, University of California Riverside; Center for Conservation Biology, University of California Riverside; Department of Plant Pathology and Microbiology, University of California Riverside; Division of Biomedical Sciences, School of Medicine, University of California Riverside

## Abstract

Air pollution poses a significant threat to the environment and human health. Most in-vivo health studies conducted regarding air pollutants, including particulate matter (PM) and gas phase pollutants, have been either through traditional medical intranasal treatment or using a tiny chamber, which limit animal activities. In this study, we designed and tested a large, whole-body, multiple animal exposure chamber with uniform dispersion and exposure stability for animal studies. The chamber simultaneously controls particle size distribution and PM mass concentration. Two different methods were used to generate aerosol suspension through either soluble material (*Alternaria* extract), liquid particle suspension (Nanosilica solution) or dry powder (silica powder). We demonstrate that the chamber system provides well controlled and characterized whole animal exposures, where dosage is by inhalation of particulate matter.

## I. INTRODUCTION

Air pollution is the presence in the outdoor atmosphere of one or more contaminants, which include particulates, gases, vapors, compounds, or biological materials in quantities, characteristics and durations that are either damaging to property or injurious to human, plant or animal life. Particulate matter (PM), including PM2.5 (particulate matter with aerodynamic diameter less than 2.5 µm) and coarser particles like dust and pollens, are major atmospheric air contaminants that continue to a pose significant threat to human health^1–3^. In recent years, extensive studies have investigated the resultant health effects of exposure to PM on human and model organisms. Numerous epidemiological studies have consistently demonstrated that air pollution is strongly associated with the morbidity and mortality from multiple cardiopulmonary and lung diseases; results indicate that PM2.5 has far more impacts on health than heretofore recognized^4–9^. One major factor is the relatively high deposition fraction of particles smaller than 2.5 microns in all regions of the lungs. The smaller the size of particulate matter, the deeper they travel into the lung; PM2.5 will even reach the alveoli. Inhaled nanoparticles can pass from the lungs into the bloodstream and extrapulmonary organs. Studies in mice have demonstrated an accumulation in the blood and liver following pulmonary exposure to a broader size range of 2∼200 nm^10^.

This chamber system allows for a route of exposure that simulates naturally occurring inhalation, as opposed to the most common method to study mouse exposures to environmental challenges. In fact, intranasal delivery may not be representative of common exposures. During an intranasal treatment, a mouse is held in a supine position while a micropipette is placed at the external nares and a concentrated solution is trickled in slowly^11,12^. Heightened concerns regarding the use of intranasal treatment, and its lack of relevance to common human exposure modes, has motivated our research aimed at exposing mice via inhalation in this study.

For better understanding the health impacts of air pollution exposure, we constructed a large-scale chamber for exposure to multiple pollutants. Featuring on whole-body exposure, this chamber houses up to 18 mice in separate cages for each experiment (^13,14^. Compared to other chronic exposure chambers featuring nose-only exposures that limits the animal activities and only allow short term exposure for each test^15,16^, our chamber frees the mice in a natural way of inhalation and delivers 7 consecutive days of exposure until the need of changing beddings or adding food supplies. Using our chamber system, we are capable of uniformly dispersing particles with sizes ranging from coarse mode particles (e.g., dust) to fine particulate matters (nano-size particles) at controlled size distributions and concentrations, while maintaining stability.

**Figure 1:**
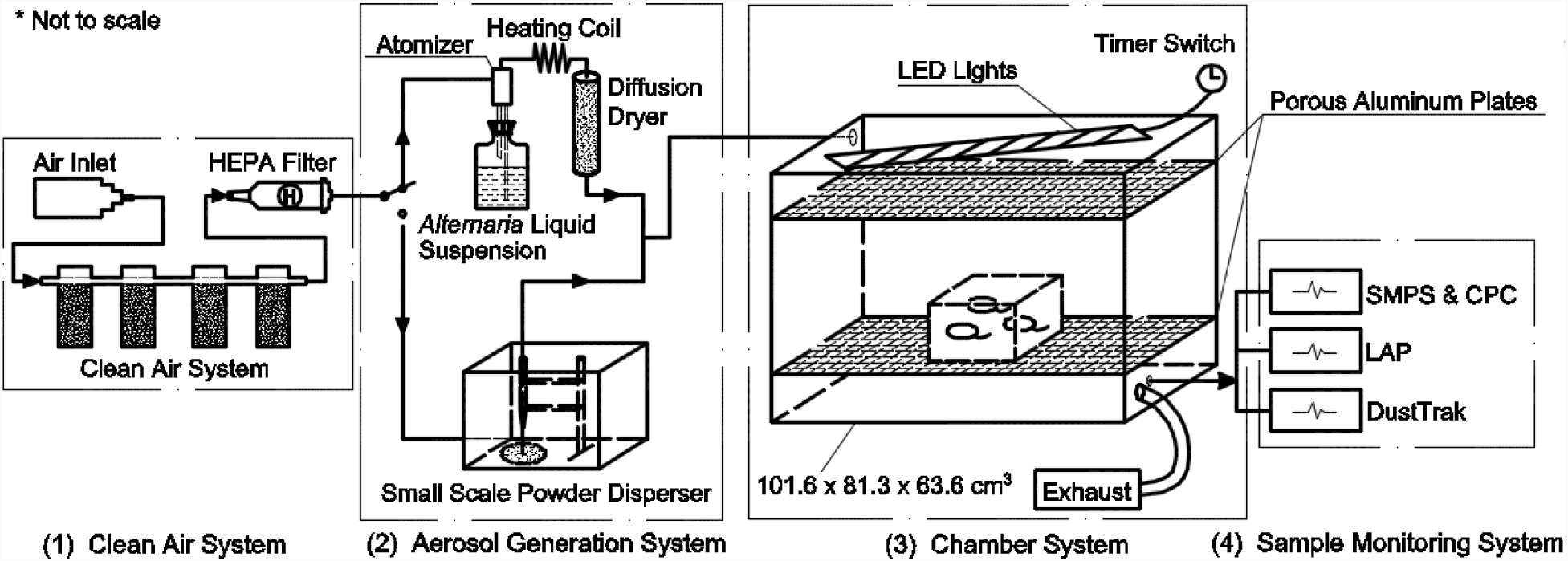
Schematic of the chamber system, including four main components: (1) Clean air system; (2) Aerosol generation system; (3) Chamber system; (4) Sample monitoring system.

## II. CHAMBER DESCRIPTION

### A. Clean air system

Lab compressed air is passed through a filter system prior to entering an Atomizer/Particle Disperser (Figure 1). The filter system consists of silica gel (absorbs water moisture), activated carbon (absorbs organics), hopcalite (absorbs CO), purafil (absorbs NOx) and a HEPA Filter (absorbs 99.97% of airborne particles).

### B. Aerosol generation system

The aerosol generation system consists of two parts, including a house-built atomizer for generating aerosol sprays from multiple liquid solutions and a Small Scale Powder Disperser for dispersing dry powder from the surface of a turntable.

### 1. Atomizer for nanoparticles

A stainless-steel atomizer generates aerosol from an ultrapure water solution of target pollutants. Compressed air passes through an orifice that creates an air jet which breaks up the solution, which is then sucked up via reduction of static pressure, generating a continuous wet aerosol spray from the solution^17^. The wet aerosol is then routed through a heated copper coil at 127 °F to transform water moisture into vapor, which is absorbed when passed through a diffuser dryer filled with indicator silica gel, replaced daily. The dried aerosols are subsequently injected into the second component, the mouse chamber. The atomizer design continuously delivers nano-size aerosol with consistent size distribution and mass concentration throughout the exposure period. Mass concentration is controlled through the concentration of aqueous solution.

### 2. Small Scale Powder Disperser (SSPD)

An SSPD from TSI (Model 3433, TSI, Minnesota, USA) was used to disperse target pollutants in powder forms, which are insoluble in water, as well as micron-size particles from 0.5 µm to 50 µm efficiently. Powders were loaded on a turntable that rotates at speeds ranging from 0.25 to 3.3 rev/hour. A stainless-steel capillary positioned just above the turntable removes powder from the surface of a turntable; the shear forces created by two flows deagglomerates the powder, which enters an expansion cone and is exhausted from the instrument and then enters the chamber. In this design, the SSPD delivers a continuous powder aerosol dose for four hours for each loading of powder on the turntable. Exposure times and mass concentrations are controlled by changing the rotation speed of the turntable.

### C. Chamber system

The mouse chamber is made of 6 clear acrylic sheets with a thickness of 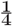 inch. The total chamber volume is approximately 540 L with external dimensions of 101.6 × 81.3 × 63.6 cm^3^ (length × width × height). Two porous aluminum plates were used to help deliver uniform dispersion by separating the aerosol inlet (upper left) and sampling ports (lower right). The large size of the chamber allows up to six mouse cages (carrying up to three mice each under normal laboratory conditions) to fit in simultaneously for exposure tests. An LED warm light string attached to a timer switch simulates a light cycle for experimental mice daily by switching on at 7 a.m. and off at 7 p.m.

To avoid light contamination, the whole chamber is covered with a customized blackout cloth and a small detachable window allows observance of mice during the exposure tests. A 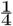 inch inlet from the upper left of the chamber was used for injection while four 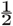 inch exhaust ports located in the lower right chamber ensured that the in-chamber pressure would not build up during aerosol injection. Another 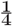 inch outlet located below the exhaust tubes was used for sampling monitoring. Before and after each test, the chamber was flushed with particle free air of at least ten chamber volumes to avoid contamination.

### D. Sample monitoring systems

Three instruments were attached to the chamber sample port to monitor the experiments.

### 1. Size Distribution and concentration measurements

Aerosol size distribution and concentration during the experiments are recorded by two different instruments. The Scanning Mobility Particle Sizer (SMPS) (Series 3080, TSI, Minnesota, USA) is used to measure particles in the range of 2 nm to 1000 nm while the Laser Aerosol Particle Size Spectrometer (LAP) (TOPAS, Germany) provides information of particle size ranging from 200 nm to 40 µm. The SMPS is widely used as a standard method for characterization of particles smaller than 1 µm in diameter with high resolution of up to 167 channels^18,19^. The LAP served as a supplemental instrument for detecting micron size particles with high resolution of up to 128 channels.

### 2. Total PM mass concentration measurements

The DustTrak^*™*^ (TSI, Minnesota, USA) Aerosol Monitor provides real-time aerosol mass readings using a light-scattering laser photometer. When equipped with different size impactors, the DustTrak measures aerosol concentrations corresponding to PM1, PM2.5 or PM10, ranging from 0.001 to 400 mg m^-3^.

## III. RESULTS AND DISCUSSIONS

### A. Mass concentration across the chamber

Figure 2 shows the PM 2.5 concentration of aerosolized ammonium nitrate (NH_4_NO_3_) solution in six different locations of the chamber plate with a standard deviation of 1.97. The DustTrak was placed inside the chamber at six locations, with the sampling port at the height of ∼10 cm to simulate mouse breathing conditions in the cages. Overall, the chamber could deliver uniform dispersion within the two porous aluminum plates that separated the aerosol inlet and the exhaust.

**Figure 2:**
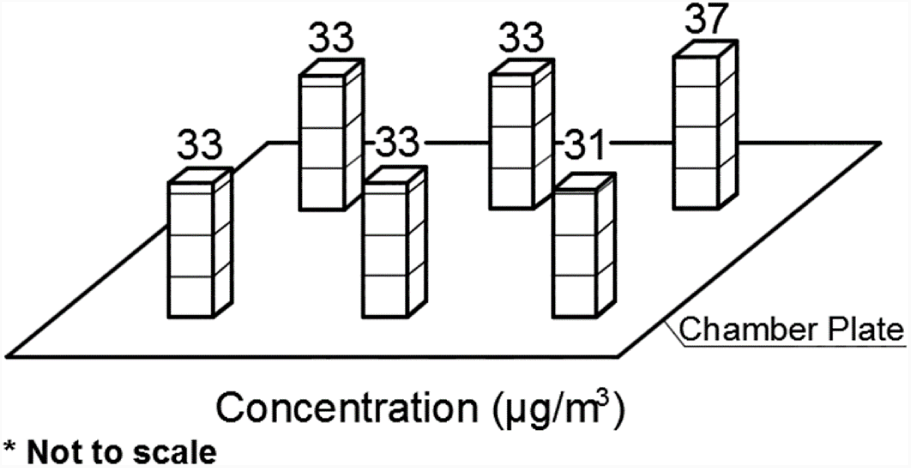
Mass concentration across the chamber. Peng et al., ASN Neuro In press, 2018

### B. Target pollutants

Multiple target pollutants, either field collected samples (swine dust/Salton Sea samples) or industrial products (Nanosilica/*Alternaria*/Silica powder), were injected into the chamber through the aerosol generation system. While the dry silica powder with the highest density (Figure 3) was dispersed using the SSPD 3433, other targets were nebulized from aqueous solutions and their size distributions can be found in Figure 5. For targets that are soluble in water or in aqueous phase, the nebulizer could provide consistent size distribution at the nanosized range, regardless of their density. All five particle suspensions in the chamber were of a size distribution to allow deep penetration into the lungs and even potential penetration into the bloodstream and extrapulmonary organs.

**Figure 3:**
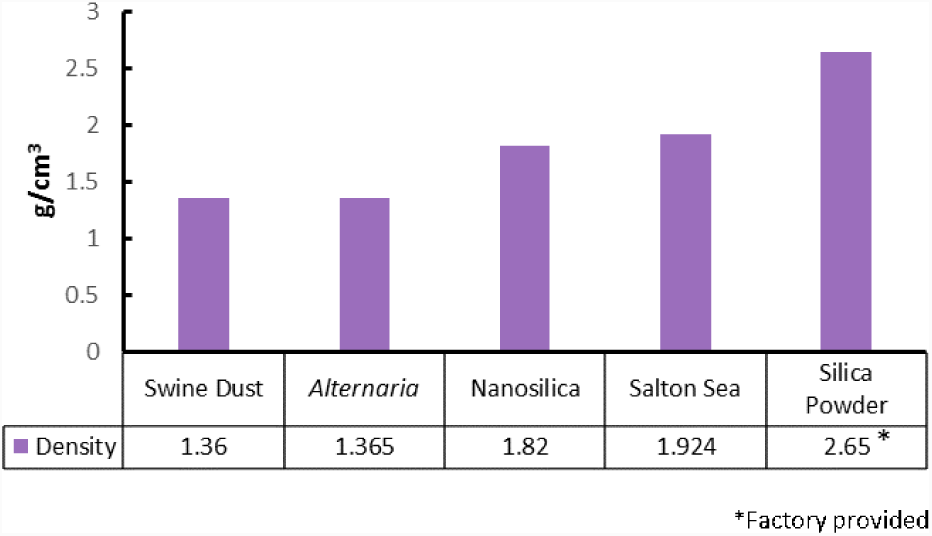
Densities of five target particle suspension in the environmental chamber. Density data was obtained using an Aerosol Particle Mass Analyzer (APM)/Scanning Mobility Particle Spectrometer (SMPS) setup, which has been modified to achieve higher transmission of particles and improved sampling frequency^20^.

**Figure 5:**
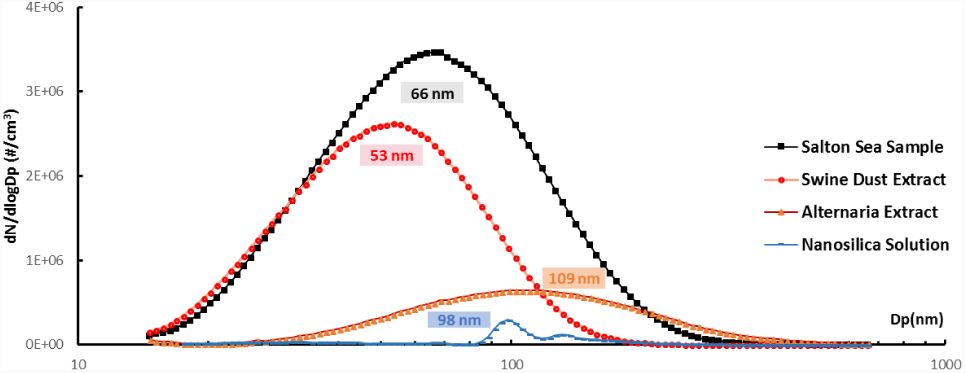
Size distribution comparison of four different nebulized target pollutants from SMPS.

### 1. Fungal extract solution

The fungus *Alternaria* alternata is a common allergen, widespread in many indoor and outdoor habitats, found to thrive on various types of vegetation. Contact with this fungus is unavoidable, given that *Alternaria* produces thousands of spores per cubic meter of air^21,22^. *Alternaria* poses a general health threat as one of the most abundant sources of aeroallergens known to trigger immune sensitization and as a primary risk factor for the development of asthma. Furthermore, exposure to *Alternaria* alternata in previously sensitized individuals is correlated with a severe increased risk of morbidity and a higher risk of fatal asthma attacks^21–24^. In children raised in desert environments, natural exposure to *Alternaria* spores induces allergic rhinitis symptoms and serves as a major allergen causing juvenile allergic asthma^21–23,25,26^. In this study, we nebulized an *Alternaria* extract solution and monitored the size distribution of *Alternaria* aerosol using the SMPS.

### 2. Salton Sea dust

Salton Sea-adjacent airborne particles we obtained by using passive collectors consisting of Teflon-coated round pans (25.4 cm in diameter) filled with quartz marbles suspended from the pan bottom with Teflon mesh, and capped with two overarching cross-braces covered in Tanglefoot to discourage roosting by birds^27^. All materials in contact with dust (i.e., pan, marbles, and Teflon mesh) were pre-washed prior to installation using first bleach, then distilled 2 M HCl, and finally distilled 3 M HNO_3_ with rinses of >18.2 M water between each reagent cleaning step. The collectors were deployed on 2.4 m tall wooden posts in open areas at each on the field sites to minimize local contributions of material from wind-suspended soil (i.e., from saltation) and nearby trees. Collectors were deployed continuously from December 2015 through March 2017 at the Dos Palmas Preserve, at 33°29’22.1”N 115°50’06.3”W. To recover dust samples from each collector in March 2017, we rinsed the marbles with >18.2 M cm water into the Teflon-coated pan, removing the marbles and mesh, then transferring the water and dust suspension to pre-cleaned 1 L LDPE Nalgene bottles. The samples were frozen and stored either at -20 or 4 °C until use. Prior to use in the chamber, samples were heated for 8h in a biosafety cabinet, in order to increase particle concentrations.

### 3. Swine facility dust

Dusts were also collected from swine concentrated animal feeding operations and prepared into a sterile aqueous extract solution^28^. Briefly, settled surface dusts were collected off horizontal surfaces within swine confinement facilities (500 ∼ 700 animals per facility). Collections were obtained from surfaces located approximately three feet above the floor of the facilities. Dusts obtained in this manner are routinely characterized for major determinants, including endotoxin and muramic acid concentrations to determine relative uniformity of samples across collection facilities and seasons. Detailed analyses of these and other determinants have been previously published^29–31^. Following collection, dust samples were placed in a buffered saline solution at 100 mg/mL concentration. Following a 1-hour incubation, samples were centrifuged to pellet all course particles and the supernatant fractions were filtered through a 0.22 micrometer pore filter. Aqueous extracts obtained following filtration were considered a 100% stock concentration and were frozen at -20 °C until use.

### 4. Nanosilica and ground silica powder

The 100 nm Non-Functionalized NanoXact^*™*^ Silica (nanoComposix Inc, California, USA) are produced via the condensation of silanes form nanoparticles that consists of an amorphous network of silicon and oxygen and the particles are monodisperse with narrow size distributions. The particles are readily suspended in polar solvents such as water and ethanol.

The MIN-U-SIL 5 GROUND SILICA powder (Western Reserve Chemical, Ohio, USA) is a natural, fine ground silica with high purity. The consistent pH and narrow size distributions allow very high loading with minimal effect on viscosity and cure rate versus synthetic silicas. The high quality, inert, white crystalline silica is available in five size distributions (5, 10, 15, 30, and 40 micron topsize).

### C. Nebulized aerosol suspension in different levels

The concentration of aerosol suspension in the chamber (Figure 4) could be controlled through changing the aqueous solution concentration, resulting in different levels of mass concentration. Three different aqueous solution concentrations of *Alternaria* extract of 0.13 g/L, 0.26 g/L and 0.52 g/L were nebulized into the chamber through an atomizer. The size distribution was acquired through the SMPS. The higher solution concentration leads to higher total number concentration in the chamber aerosol suspension without significantly changing the particle mode diameter of 100 nm.

**Figure 4:**
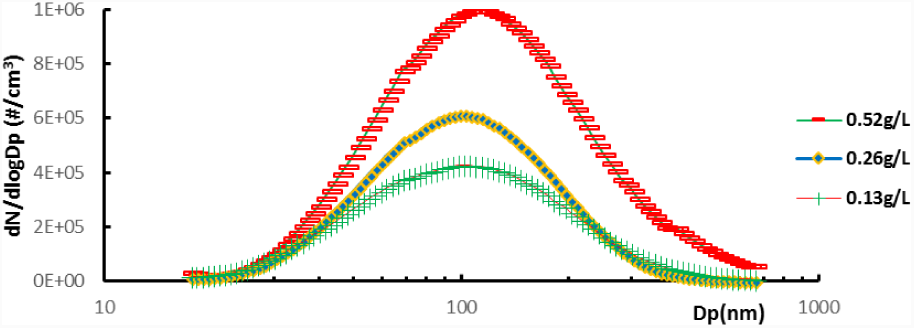
Size distribution of *Alternaria* aerosol in three different levels. The Y axis stands for particle number concentration dN/dlogDp (#/*cm*^3^) while the X axis represents particle size on a logarithmic scale.

Four different types of target pollutants were nebulized into the chamber at a controlled number/mass concentration (Figure 5 Figure 6). While the mode diameters of these aerosol range from 53 nm to 109 nm, all of them can pass from the lungs into the bloodstream and extra-pulmonary organs. Higher number concentration does not always result in higher mass concentration since larger size particles dominate the total mass concentration. For example, the swine dust extract had a much lower total mass concentration than that of the *Alternaria* extract (Figure 6) even though it had significantly higher number concentration (Figure 5).

**Figure 6:**
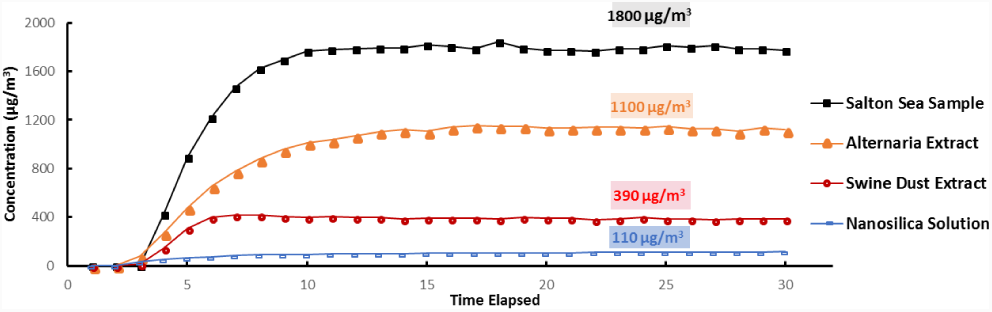
Consistent mass concentration of target pollutants over time.

Mass concentration could be controlled from the micro gram per cubic meter range to milligram per cubic meter range (Figure 6). The chamber was saturated with continuous aerosol injection and remain consistent throughout the whole experiment, providing a much more effective and protracted dose delivery than the intranasal method.

### D. Powder-silica aerosol from SSPD 3433

Occupational exposures in different trades range from the lowest on operation engineers (0.075∼0.720 mg m^-3^) to the highest on painters (1.28∼13.5 mg m^-3^)^32^, which significantly exceeds the US Occupational Exposure Limit (OEL) of 0.05 mg m^-3^ for respirable silica^33^.

To simulate exposures to respirable silica among construction sites in the USA, two 4-hour silica doses were injected into the chamber each day (Figure 7). The first dose was given from 8 a.m. to 12 p.m. while the second dose was given from 4 p.m. to 8 p.m. Figure 7 shows ten doses injected over 5 days with average dose concentration of around 5mgm-3 and the overall average concentration was 1.43 mg m^-3^. Figure 8 shows the size distribution of dispersed silica powder with a mode of around 306 nm.

**Figure 7:**
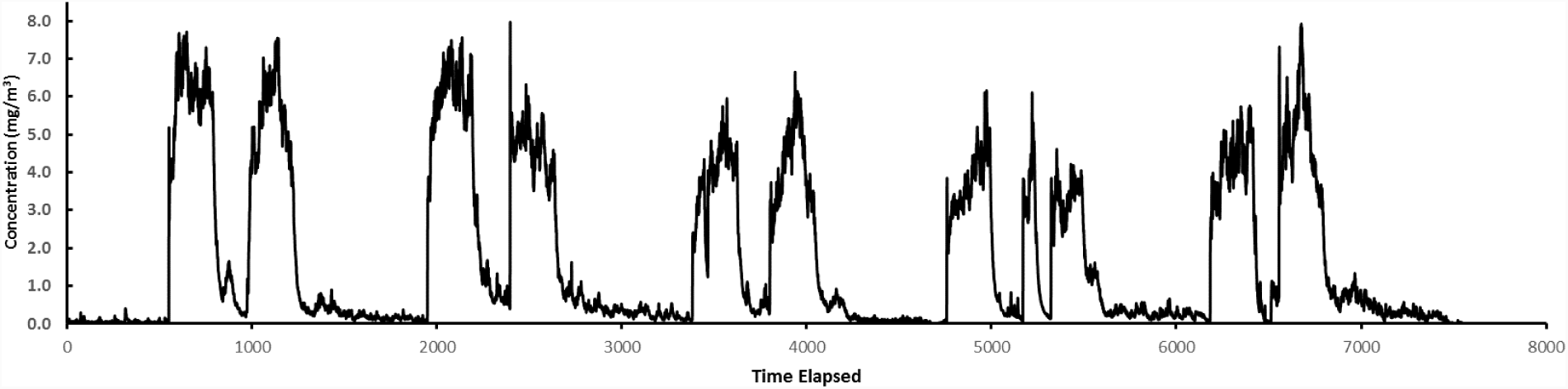
Mass concentration of powder silica aerosol throughout the five-day exposure.

**Figure 8:**
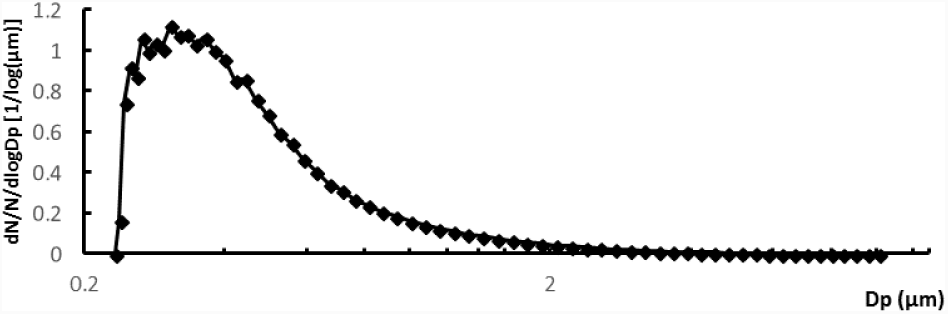
Size distribution of powder silica aerosol. The vertical axis stands for particle number distribution dN/N/dlogDp [1/µm] while the horizontal axis represents particle size on a logarithmic scale.

## IV. CONCLUSION

Here we developed a large-scale multipurpose animal exposure chamber for health effects investigation of air pollutants. Compared to other chamber systems featuring nose-only or activity-limited whole-body exposure, this system provides continuous exposure through the natural inhalation route at controlled concentration, size distribution and duration for both chronic, sub-chronic and acute epidemiological studies. Our method is also easily reproducible and has the potential to mimic the real atmospheric environment by adding multiple air contaminants at the same time for health investigations.

## ACKNOWLEDGMENTS

These studies were funded by grants from UCR Office of Research Seed grant and the UCR Office of Research for funding, as well as NSF EAR-1744089. We would like to thank Chen Le for the chamber system schematic design and Chelsea Carey, Taryn Barsotti, and Geoffrey Logan who assisted with collector set-up or processing of the samples in the lab.

